# Fractal and Machine Learning analyses of MALDI-TOF Mass Spectrometry data in glioblastoma

**DOI:** 10.1101/2025.03.03.641183

**Authors:** Lucas C. Lazari, Ghasem Azemi, Carlo Russo, Livia Rosa-Fernandes, Suely K. N. Marie, Antonio Di Ieva, Giuseppe Palmisano

## Abstract

Data pre-processing is a critical step in the analysis of MALDI-TOF MS spectra for machine learning applications, typically involving steps such as spectra trimming, baseline correction, smoothing, transformation, and peak picking or spectral binning. While traditional approaches focus on protein/peptide peaks as features, this study explores a novel method of feature extraction by treating MALDI-TOF spectra as time-series data. This study investigates the use of computational fractal-based analysis to assess the complexity of MALDI-TOF spectra. Fractal analysis, previously successful in glioblastoma diagnosis using MRI, was applied here to proteomics data for the first time. By treating each MALDI spectrum as a time series and calculating its fractal dimension using various algorithms, machine learning models were trained to differentiate between glioblastoma patients and healthy controls. We demonstrate that fractals are sufficient to obtain accurate models for glioblastoma diagnosis, despite still underperforming when compared to the traditional feature extraction method. We also show that fractals can be used as support features to increase model performance. This work highlights the potential and limitations of fractal analysis in proteomics, offering a new perspective for disease diagnosis and broadening the applicability of time-series data analysis in mass spectrometry.

## 1. Introduction

Data pre-processing is an essential step for analysis with MALDI-TOF MS spectra for machine learning applications. The pre-processing steps usually consist in spectra trimming, baseline correction, smoothing, transformation and peak picking ^1^. Conversely, the peak picking step can be replaced by spectral binning, in this way the features are no longer protein/peptides m/z peaks, rather the whole spectra ^2^. In the previous chapters we demonstrated how the features (proteins/peptides peaks with a given intensity) generated by these approaches can be used to train machine learning models. Now we demonstrate a new approach to extract features from MALDI-TOF data by using time-series data analysis.

The main question was whether or not a MALDI spectrum could be considered a time-series data. Although the spectrum mass-to-charge ratio could be converted to time, since it is calculated based on the time-of-flight of the molecule before it reaches the analyzer, the proteomics itself does not reflect a temporal data, meaning that the spectrum of a determined sample was acquired in a given time (i.e., the moment of serum collection). Therefore, for proteomics data to be considered truly time-series, a collection of spectra should be retrieved from different samples that were collected at different times. With this limitation in mind, we decided test the limits of what can be done with a MALDI-TOF spectra and test the capabilities of time-series data analysis applied to mass spectrometry.

In this context, working with the spectrum as a time-series allows the use of different data analysis techniques. Among these techniques, computational fractal-based and analysis provides potential useful markers for the assessment of time series’ complexity.

One of the most used parameters in fractal geometry is the fractal dimension (FD), used to characterize the self-similarity and repeating patterns across different scales through non-Euclidean dimensions of an object ^3^. The fractal dimension of an object increases with its structural complexity ^4^. Fractal analysis has already been successfully employed in glioblastoma diagnosis and prognosis using imaging techniques such as MRI, demonstrating that high grade tumors tend to have structural and angioarchitectural higher fractal dimension values than normal brain or lower grade tumors ^5–7^. Fractal analysis can also be applied to time series data, too, where the time series pattern may exhibit fractal scaling properties ^8^. Regarding proteomics data, there is no report in the literature involving the use of fractals.

Due to the lack of research involving fractal dimension in mass spectrometry, we decided to explore this topic using MALDI-TOF MS data in glioblastoma diagnosis. For that, each MALDI spectrum was treated as a time series, and its fractal dimension were calculated using different algorithms. These measurements were then used to train machine learning models to distinguish between glioblastoma patients and healthy controls. We demonstrate the potentials and limitations of this approach in proteomics, highlighting its application in different diseases.

## 2. Methods

In this study, we evaluated the utility of fractals for classifying GBM patients by comparing models trained with fractal dimensions to models trained with binned spectra. For classification purposes, we used the entire spectra of the samples. The binning procedure was intended to reduce the effect of inter-sample variability (such as peak shifts) and to standardize all samples to the same dimensionality. However, in our case, all samples already had the same dimensionality. The model training was performed in a computer with an AMD Ryzen 5 5600 CPU. A graphical abstract about the methods employed in this chapter is present in Figure 22.

### 2.1. Dataset overview

The MALDI-TOF MS spectra from glioblastoma and control patients were obtained from PRIDE database (PXD060922). A total of 283 files were retrieved from the database. The downloaded files included 211 oncological patients, 62.44.6% of whom were male and 37.56% female, with ages ranging from 0.5 to 69 years. The majority of patients (78.68%) were diagnosed with GBM, while 11.17% had grade 2 (AG2) tumors, and 10.15% had grade 3 (AG3) tumors. The control group consisted of 72 individuals, 65.28 % female and 34.72% male, with ages ranging from 28 to 86 years (Table 7).

**Table 1.** Cohort summary, describing the distribution of age, sex and tumor grade of the patients involved in this study.

|  | Age (years) | Sex | Grade |
| --- | --- | --- | --- |
| ***Oncological Patients*** | Min - 0.5 | F - 37.56% | GBM - 78.68% |
|  | Max - 69 | M - 62.44% | AG2 - 11.17% |
|  | Mean - 49.77 |  | AG3 - 10.15% |
| ***Controls*** | Min - 28 | F - 65.28% | - |
|  | Max - 86 | M - 34.72% | - |
|  | Mean - 56.19 |  | - |

### 2.2. Spectra files acquisition and preprocessing

The mzML files were pre-processed in R using the MALDIQuant package 167. All spectra were first smoothed using the “SavitzkyGolay” method with a half window size of 10. Baseline correction was then performed with the TopHat method, and finally, the intensities were normalized to the total ion current. Although all spectra were processed concurrently, each spectrum underwent the preprocessing procedures individually, ensuring that no data leakage occurred during the preprocessing step.

### 2.3. Fractal dimension calculation

The Petrosian, Katz, Higuchi and Detrended fluctuation FDs were calculated using the AntroPy library ^9^. The MALDI-TOF MS spectra was treated as a time-series data, using the intensities from 2000 m/z to 20000 m/z. The fractal dimension values were calculated using the complete spectra and the spectra divided in windows of fixed size (600, 800, 1000, 1200, 1400 and 1600 delta m/z values), meaning that the intensity vector was divided in intervals of fixed size and all fractals’ dimensions were calculated within that interval. This approach generates more features as the window size decreases. All parameters from the AntroPy package were set to their default values (higuchi kmax = 10).

### 2.4. Spectra binning

Using the method described in Weis et al. (2022), the binning procedure was performed and the pre-processed spectra were partitioned into non-overlapping bins of 3 m/z. The intensities within each bin were summed ^2^. This procedure not only standardize the data but also greatly reduces the dimensionality of the spectrum.

### 2.5. Machine learning for classification

The procedure for tumour classification using fractals and the binned spectra followed the same pipeline. First, the dataset was split into five folds of train and test sets using the stratified k-fold function of sklearn python library, this was done to ensure that both approaches were trained using the same set of samples in each fold, thus ensuring that the results are comparable. Then, a 5-fold nested-repeated 3-fold cross-validation was performed for hyperparameter tunning and model evaluation. The pipeline used in this work for model training is showed in Figure 23.

After selecting the best set of hyperparameters in the inner loop, the best model is evaluated using the test set, this is performed across all the dataset divided in 5 folds. The models’ performances were evaluated for the dataset with and without oversampling and with different methods for feature selection (BorutaPy ^10^, Principal Component Analysis (PCA), Partial Least Squares (PLS) and Receiver-operating Characteristic Curve (ROC) Area Under the Curve (AUC)). The pipeline with highest performance that was used to generate all data in the results section was without Synthetic Minority Over-sampling Technique (SMOTE) to oversample the control class, scaling the training and test sets using the StandardScaler function from sklearn (the scaler was fit in the training set) and BorutaPy for feature selection (n_estimators = “auto”, alpha = 0.001), these procedures were performed within each fold. The resulting features were used to train three models (Random Forest, LightGBM and SVM), and performance metrics including sensitivity, specificity and balanced accuracy were evaluated. Finally, the best model was selected to run another 5-fold cross validation with Shapley Additive Explanations (SHAP library) to evaluate the feature contribution, for that we used the KernelExplainer algorithm. The performance metrics for all other feature selection methods and the models with oversampling are available in the Supplementary Table 1.

Confusion matrices were created to calculate the performance metrics for each test performed, ROC and PR curves were also generated. A mean spectrum was generated for each group and the interquartile range was calculated and plotted as a shaded area in the mean spectrum. The most important features accordingly to SHAP^11^ for both fractals and binned features were plotted alongside the spectrum to search for regions of interest.

## 3. Results

### 3.1. Classification performance in varying window size

To find the optimal window size for the classification task using fractals based on model performance (balanced accuracy), we segmented the intensities arrays in different sizes (600, 800, 1000, 1200, 1400 and 1600 delta m/z values) and also tested the classification performance using the complete intensity array from the entire m/z range. Therefore, we searched for the hyperparameters set and trained the three models on each window size and compared the balanced accuracy, sensitivity and specificity. The results demonstrated that the window size of 1000 achieved the highest balanced accuracy (90.2%) using a SVM model (Figure 3A). The lowest performance was obtained without spectrum segmentation. All models’ performances for this step are reported in the Supplementary Table 1.

**Figure 1:**
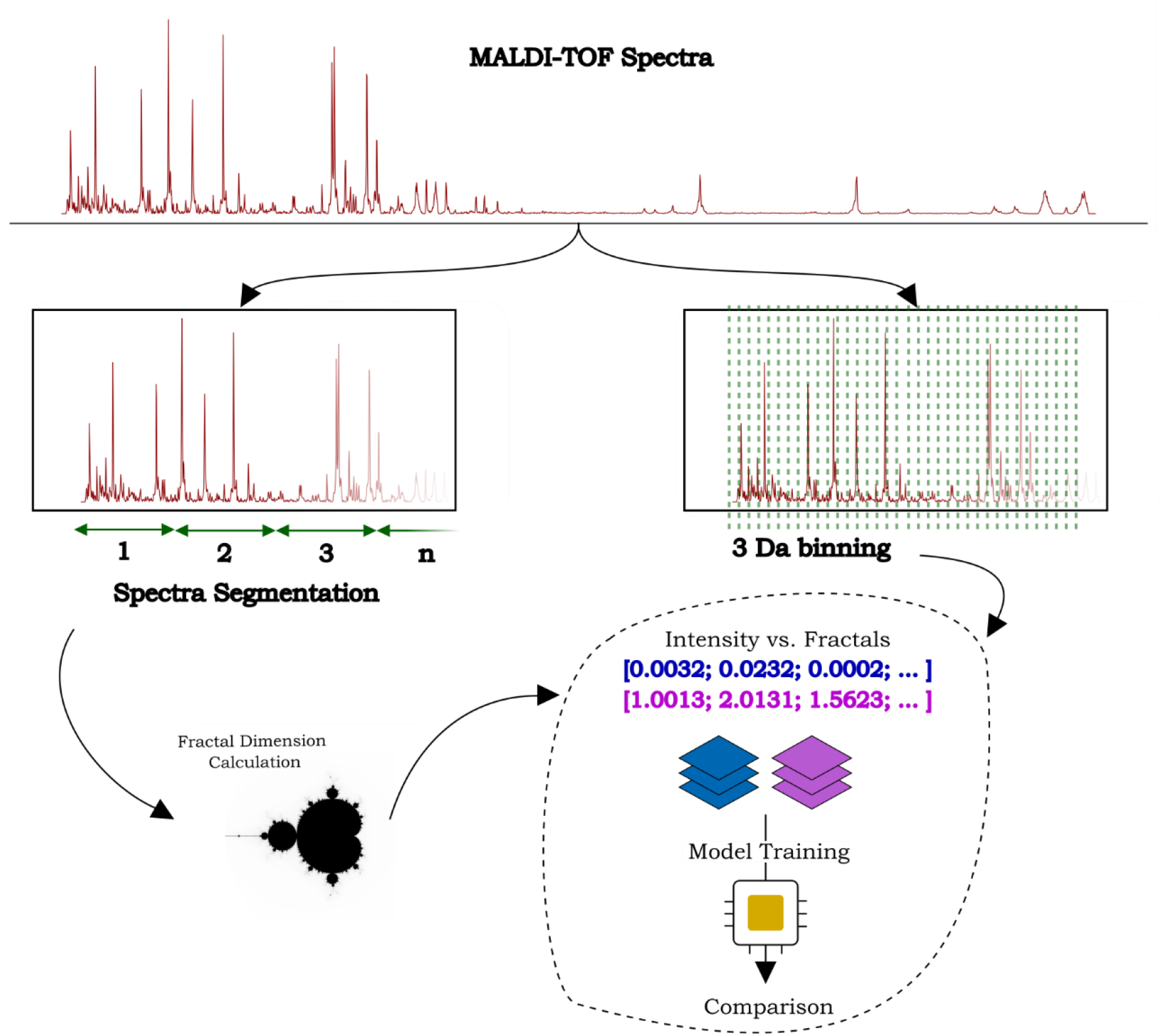
Representation of the methods to perform fractal analysis in mass spectrometry data. The methodology consists of two approaches, a classical approach to generate features by using the whole binned spectra, and a newly developed method based on features generated by calculating the fractal dimensions of the segmented spectra.

**Figure 2:**
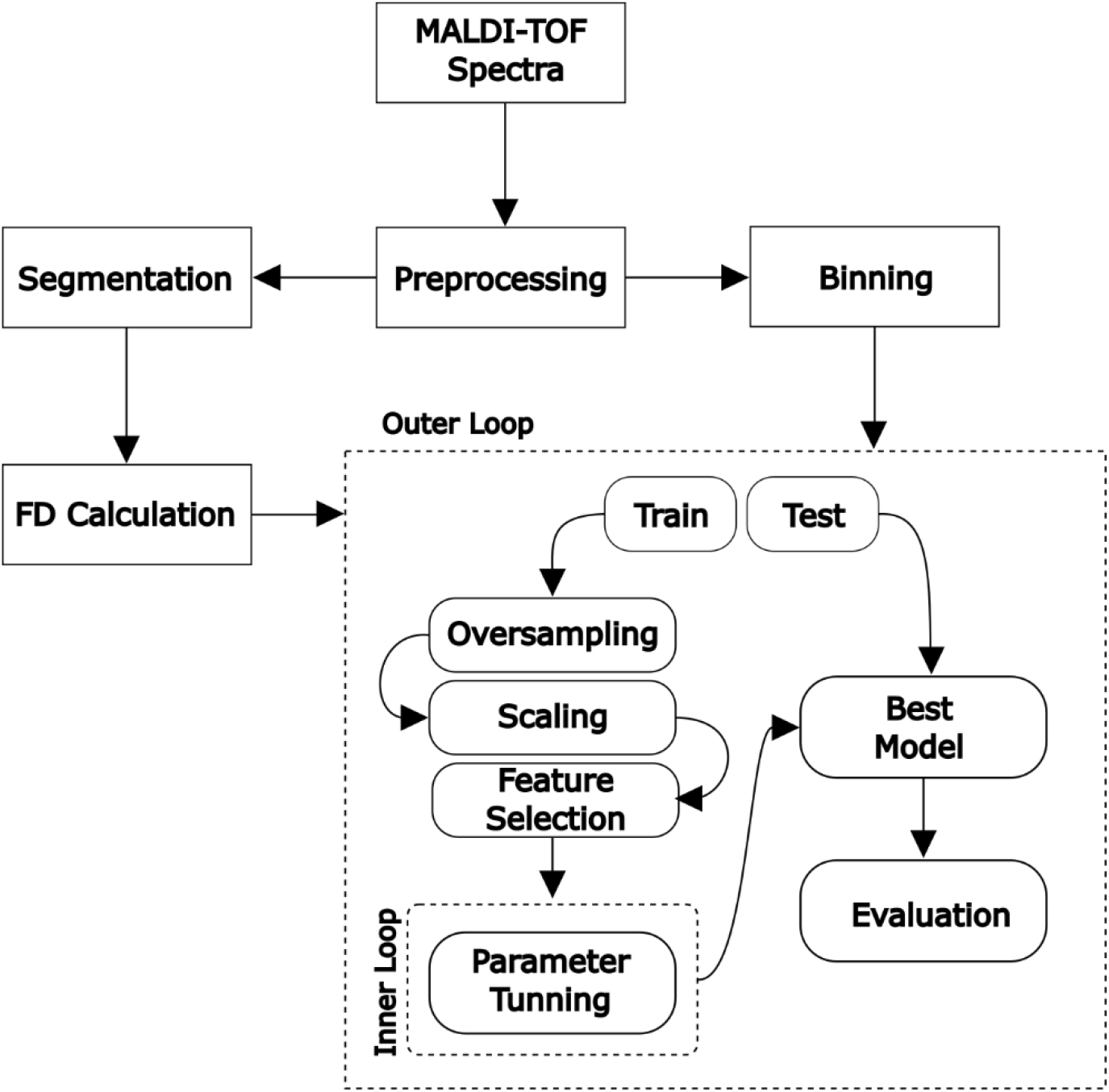
Pipeline used for model training and evaluation of fractals in mass spectrometry. An outer loop is used to separate the date into train and test sets. Then, the training set is processed and used in an inner loop for hyperparameter tunning through a 3-Fold Cross-Validation. The best model is then selected and evaluated using the hold-out test set. The scaling and feature selection procedures are performed only in the training set, then passed to the test set to avoid over fitting.

**Figure 3:**
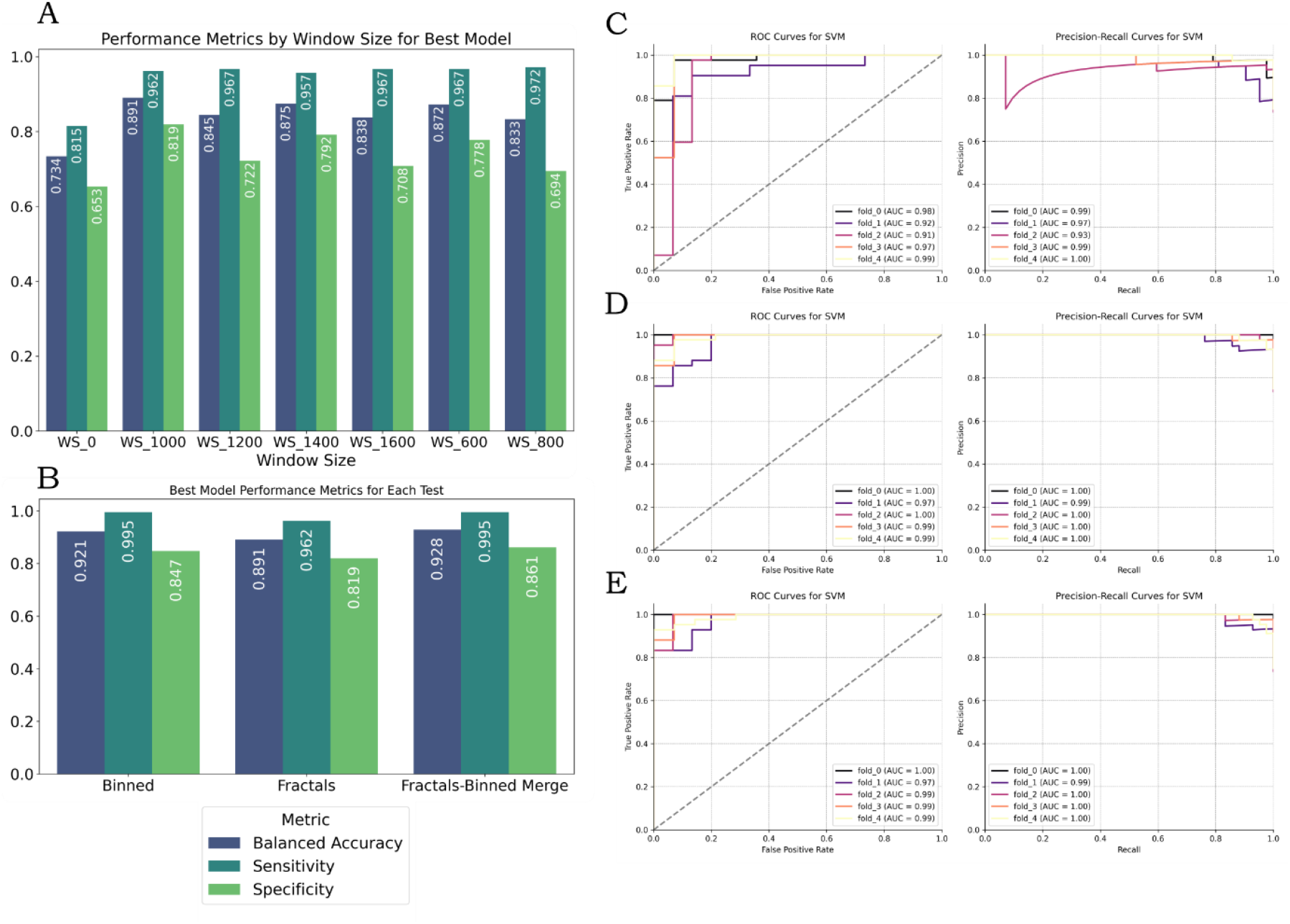
Performance of the models trained with fractals and binned data. (A) Performance metrics for the best model in each window size tested. (B) Performance metrics for the best model in each dataset tested. (C) ROC and PR curves for the best model trained with the fractals features dataset. (D) ROC and PR curves for the best model

### 3.2. Comparison of fractals and binned spectra for classification

From a total of 32 intervals using the optimal window size of 1000, the total number of fractals features were 128 prior to feature selection (these features were generated by dividing the intensity array into sub-arrays of size 1000 and calculating the FD dimensions within that sub-array), which were reduced to 73 by BorutaPy. The total analysis time, from fractals dimensions calculation to the five-fold cross-validation with optimized parameters took approximately 4 minutes. As described in the previous section, the highest performance obtained using fractals was obtained by the SVM model, achieving a balanced accuracy, sensitivity and specificity of 89.1%, 96.2% and 81.9% respectively (Figure 24B). For each fold we used SHAP to evaluate the most important features for the classification task and how they impacted the model’s decision making (Figure 4). We then selected the top 10 features across all folds and searched the intensity intervals in which they were calculated. This was performed to find the discriminating regions in the spectra, which could be used to detect potential biomarkers (Figure 5). The output provided by SHAP demonstrated that different fractals dimensions calculation methods in the same region of the spectra presents different behaviors. For instance, Detrended FD at the region of number 7 have a negative correlation with the classification of the positive class (GBM patients), meaning that when this value is high, the model tends to classify the individual as negative class (control). However, Katz FD in the same region had the opposite behavior. This strengthen the idea of using multiple FD calculation methods, being beneficial to capture the complex nature of MALDI-TOF MS proteomics. Interestingly, most of the fractals features across all folds had a clear impact for the model’s decision making, meaning that it is simpler to interpret whether the increase or decrease of a feature will impact the model’s classification of the positive class when compared to the binned spectrum (Figure 4).

**Figure 4:**
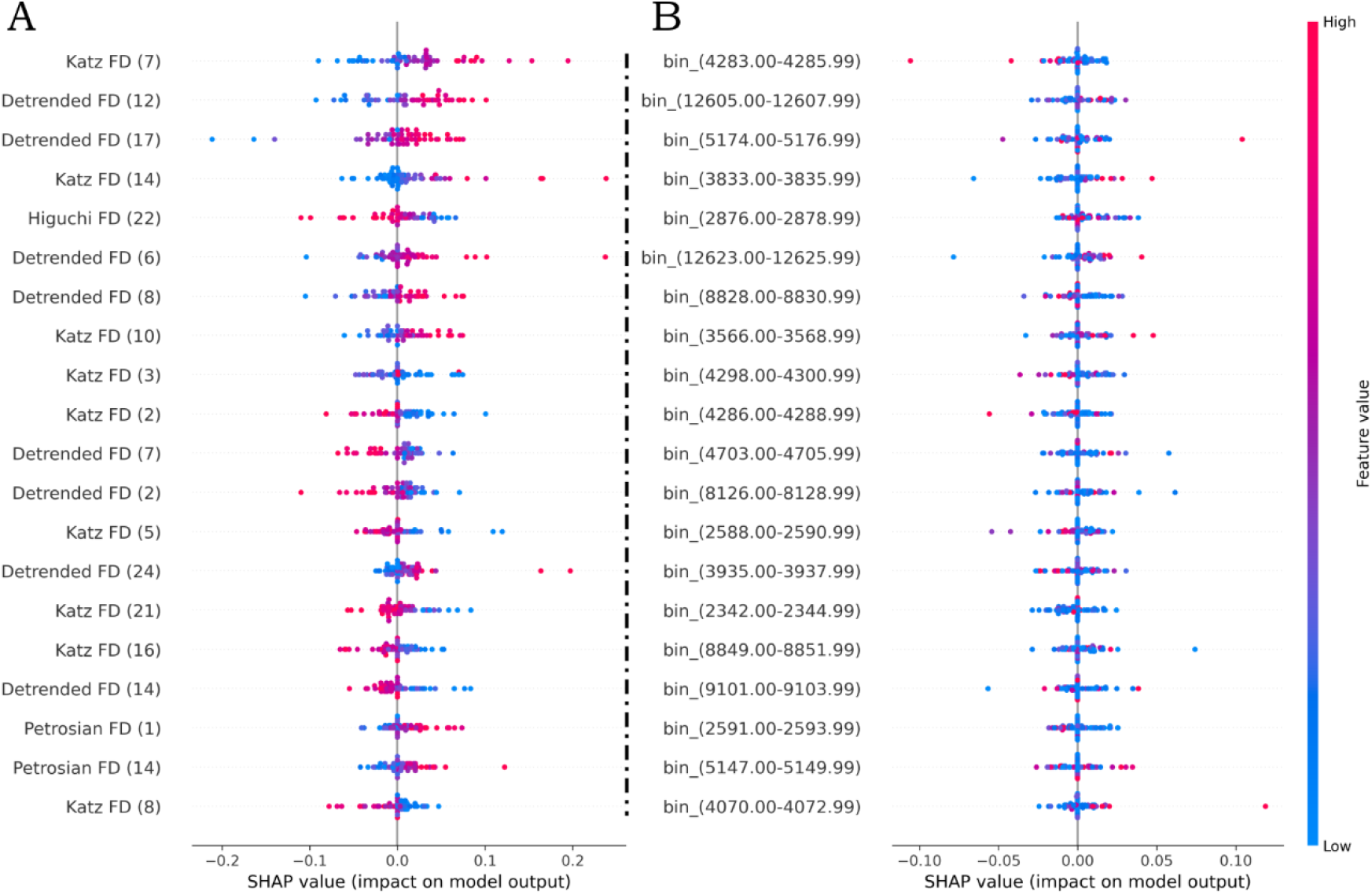
SHAP results for model explanation. A) SHAP output for the first fold of the fractals features dataset. (B) SHAP output for the first fold of the binned features dataset. SHAP value indicates the impact that feature has on model output, positive values imply an impact in the classification of the positive class, while negative values imply the opposite. The color map indicates how the feature impacts on model decision, e.g., if the feature has a high value and a high SHAP value, the increase of this feature is a characteristic of the positive class.

**Figure 5:**
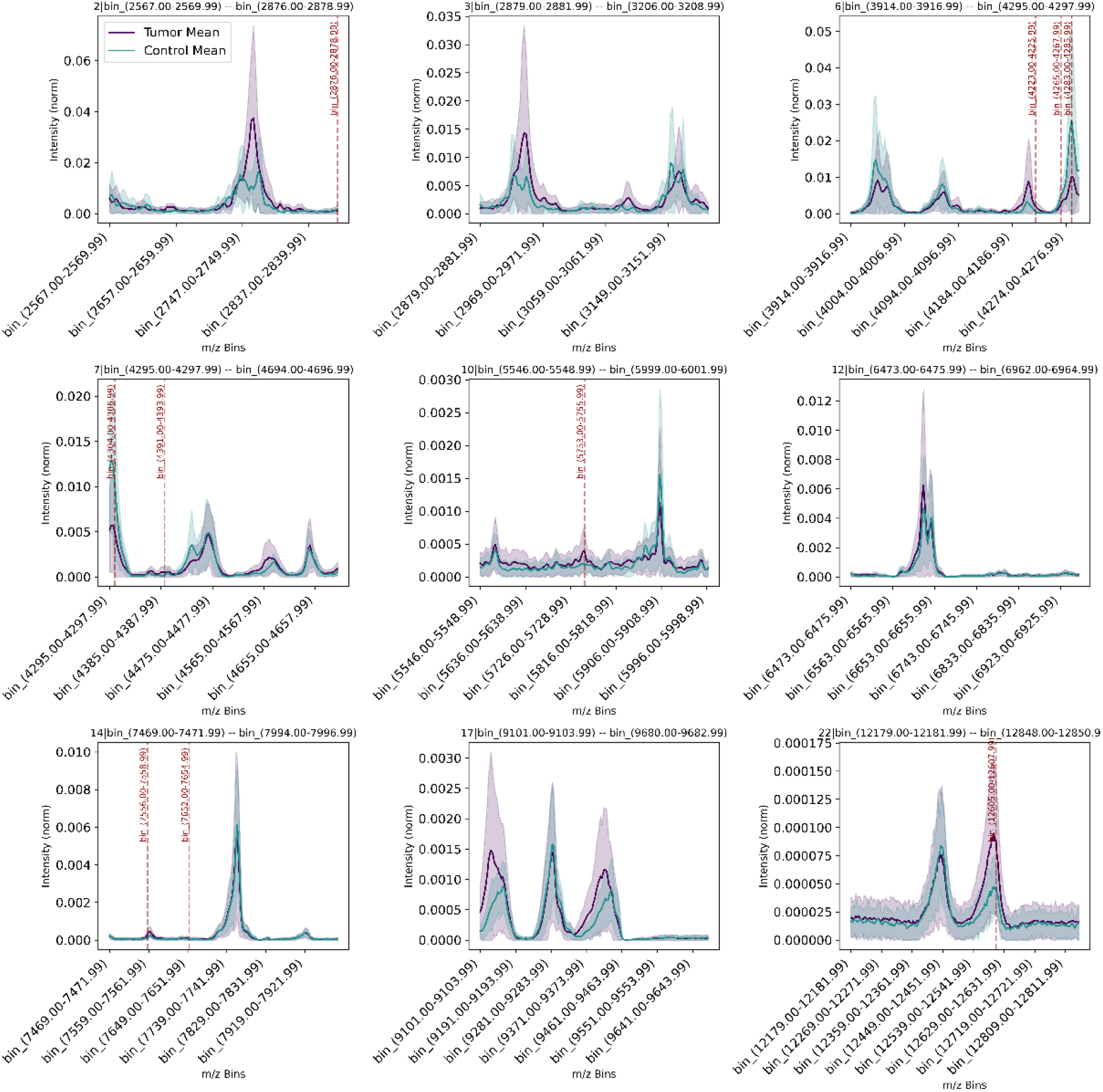
Zoomed MALDI-TOF MS spectra in the most important regions according to SHAP. On top of each plot, the first number is related to the window that is being shown, followed by relevant bins within that window. The shaded area is the interquartile range, the dashed red lines are relevant bins accordingly to SHAP.

For the binned spectra, the number of features prior feature selection was 4336 (generated by grouping the m/z into 3 Da intervals and summing their intensities), which were reduced to 470 using BorutaPy. The total analysis time in this case was approximately 16 minutes, however, the performance was higher than the fractals dataset. Using the binned spectra, we achieved a balanced accuracy, sensitivity and specificity of 93.0%, 98.6% and 87.5% respectively (Figure 3). We applied SHAP in all folds to find the most important bins and to evaluate how they impact the model’s decision making. Conversely from fractals features, the binned features across all folds are more concentrated at the centre of SHAP value, meaning that is difficult to evaluate how the increase or decrease of a feature will impact the model’s classification of the positive class (Figure 4B). The binned features that were present in the same region as the selected fractal features were pointed in the plot (Figure 5).

All models’ performances, their respective ROC and PR curves, and the best hyperparameters sets are available in the Supplementary Material 1. The SHAP output for each fold for both fractals and binned features is available in the Supplementary Material 1.

Following SHAP results, we extracted the top five features of each fold and selected the unique features as candidates for biomarkers. Figure 5 shows the unique spectra regions that contained the top five fractals features and the top five bins that fall within that region. This demonstrates that fractals can be used to tune up the search for relevant regions that could be further investigated using other proteomics techniques, with the final aim to identify proteins which might be useful as biomarkers.

The regions 2, 3 and 7 show a clear distinction between the groups, with the appearance of mass peaks with different intensities between the groups.

## 4. Discussion

Fractals have found significant applications in the analysis of time series data in a wide range of fields, such as climate and medicine research ^12,13^. A MALDI-TOF MS spectrum is not typically considered time series data, as time series data typically consist of sequential measurements taken at specific time interval, while MALDI-TOF spectrum represents data as intensity versus m/z ratio, reflecting the distribution of molecular masses at a single point in time, lacking the temporal sequence characteristic to time-series data. Apart from one work that modelled MALDI-TOF MS data as time series from spectral clustering ^14^, no other work has analyzed it directly as a time series data. In our study, we investigated whether the fractal analysis of time series data could be directly applied to MALDI-TOF MS data.

In the present study we re-analyzed the MALDI-TOF MS data from a GBM cohort to develop a new data processing pipeline for machine learning classification using fractal analysis. Although the use of MALDI-TOF MS data has already been employed in this thesis and has been explored in the literature^15^, this work demonstrated for the first time that MALDI-TOF MS spectrum has fractals properties and the fractal dimensions of the protein profile of healthy and GBM patients differ, suggesting the existence of a GBM’s spectroscopic fractal fingerprint. The use of fractals as features for model training yielded a high performance in classification, but lower when compared to the binned features. However, the merge of both feature types caused a slightly increase in overall model performance. This indicates that although fractals do not outperform spectrum binning, fractal analysis can be still used as a complementary analytic tool. A clear advantage to use fractals in this case is to reduce processing time, since fractal analysis was significantly faster than spectrum binning (approximately 4 times faster), being a promising technique for the analysis of very large datasets.

SHAP analysis demonstrated that fractals can also be used to search biomarkers by finding spectra regions with highly-significant features. These regions of interest could be evaluated using other proteomics techniques to identify proteins in that mass range and assessing their potential as biomarkers.

The possibility to use fractal analysis in mass-spectrometry data opens a new field for proteomics, where fractals can be explored not only in GBM research, but in many other diseases. To this point, we found that fractals dimensions can be used to complement the dataset to enhance model performance, with more investigation needed to assess other possible applications of this technique in proteomics. Also, this work demonstrated that the MALDI-TOF MS spectrum can be used as a time series data, which allows researchers to explore a wide range of time series analytical tools.

## 5. Conclusions

This study demonstrates that fractal dimensions of MALDI-TOF MS spectrum can be used to characterize glioblastoma in liquid biopsies. Fractals can also be used to search for regions of interest in the spectra, that could be further translated into biomarker discovery. Although it could not outperform the models trained with binned spectra, fractals can serve as a complement to increase model performance. Since the analysis time was significantly lower, fractals could be an alternative to deal with very large datasets. This is the first report of fractals being associated with mass spectrometry data and it opens a new field for research in proteomics. The methodology employed in this work could be used not only to study glioblastoma, but many other diseases that are already being explored using MALDI-TOF data. It also opens the possibility to test fractal dimensions in other mass spectrometry data types.

