## Supplementary Materials for "Fractal and Machine Learning analyses of MALDI-TOF Mass Spectrometry data in glioblastoma"

### Supplementary Material

**Table 1:** All models performances for each category tested.

| *Category* | *Model* | *Mean Sensitivity* | *Mean Specificity* | *Mean Balanced Accuracy* |
|---|---|---|---|---|
| *Fractals* | RandomForest | 0.976303318 | 0.708333333 | 0.84231833 |
| | LightGBM | 0.971563981 | 0.722222222 | 0.8468931 |
| | SVM | 0.962085308 | 0.819444444 | 0.89076488 |
| *Binned* | RandomForest | 0.976303318 | 0.819444444 | 0.89787388 |
| | LightGBM | 0.966824645 | 0.833333333 | 0.90007899 |
| | SVM | 0.995260664 | 0.847222222 | 0.92124144 |
| *Fractals-Binned Merge* | RandomForest | 0.981042654 | 0.819444444 | 0.90024355 |
| | LightGBM | 0.976303318 | 0.833333333 | 0.90481833 |
| | SVM | 0.995260664 | 0.861111111 | 0.92818589 |

**Table 2:** Hyperparameters selected by random search in each fold for each model and category:

| FOLD | MODEL | PARAMETERS | CATEGORY |
|---|---|---|---|
| **FOLD_0** | RandomForest | {'max_depth': 44, 'n_estimators': 393} | Fractals |
| | RandomForest | {'max_depth': 44, 'n_estimators': 448} | Binned |
| | RandomForest | {'max_depth': 44, 'n_estimators': 448} | Merged |
| | LightGBM | {'learning_rate': 0.20475110376829186, 'n_estimators': 370, 'num_leaves': 32} | Fractals |
| | LightGBM | {'learning_rate': 0.12973169683940733, 'n_estimators': 202, 'num_leaves': 48} | Binned |
| | LightGBM | {'learning_rate': 0.0849080237694725, 'n_estimators': 370, 'num_leaves': 37} | Merged |
| | SVM | {'C': 37.55401188473625, 'gamma': 'scale'} | Fractals |
| | SVM | {'C': 37.55401188473625, 'gamma': 'scale'} | Binned |
| | SVM | {'C': 37.55401188473625, 'gamma': 'scale'} | Merged |
| **FOLD_1** | RandomForest | {'max_depth': 44, 'n_estimators': 448} | Fractals |
| | RandomForest | {'max_depth': 44, 'n_estimators': 120} | Binned |
| | RandomForest | {'max_depth': 44, 'n_estimators': 171} | Merged |

| | | | |
|---|---|---|---|
| | LightGBM | {'learning_rate': 0.0849080237694725, 'n_estimators': 370, 'num_leaves': 37} | Fractals |
| | LightGBM | {'learning_rate': 0.18198808134726413, 'n_estimators': 234, 'num_leaves': 50} | Binned |
| | LightGBM | {'learning_rate': 0.04636499344142013, 'n_estimators': 376, 'num_leaves': 62} | Merged |
| | SVM | {'C': 37.55401188473625, 'gamma': 'scale'} | Fractals |
| | SVM | {'C': 37.55401188473625, 'gamma': 'scale'} | Binned |
| | SVM | {'C': 37.55401188473625, 'gamma': 'scale'} | Merged |
| **FOLD_2** | RandomForest | {'max_depth': 35, 'n_estimators': 291} | Fractals |
| | RandomForest | {'max_depth': 44, 'n_estimators': 448} | Binned |
| | RandomForest | {'max_depth': 44, 'n_estimators': 393} | Merged |
| | LightGBM | {'learning_rate': 0.0849080237694725, 'n_estimators': 370, 'num_leaves': 37} | Fractals |
| | LightGBM | {'learning_rate': 0.02999498316360058, 'n_estimators': 187, 'num_leaves': 65} | Binned |
| | LightGBM | {'learning_rate': 0.0849080237694725, 'n_estimators': 370, 'num_leaves': 37} | Merged |
| | SVM | {'C': 37.55401188473625, 'gamma': 'scale'} | Fractals |
| | SVM | {'C': 37.55401188473625, 'gamma': 'scale'} | Binned |
| | SVM | {'C': 37.55401188473625, 'gamma': 'scale'} | Merged |
| **FOLD_3** | RandomForest | {'max_depth': 44, 'n_estimators': 199} | Fractals |
| | RandomForest | {'max_depth': 35, 'n_estimators': 314} | Binned |
| | RandomForest | {'max_depth': 44, 'n_estimators': 187} | Merged |
| | LightGBM | {'learning_rate': 0.12973169683940733, 'n_estimators': 202, 'num_leaves': 48} | Fractals |
| | LightGBM | {'learning_rate': 0.20475110376829186, 'n_estimators': 370, 'num_leaves': 32} | Binned |
| | LightGBM | {'learning_rate': 0.18198808134726413, 'n_estimators': 234, 'num_leaves': 50} | Merged |
| | SVM | {'C': 18.44347898661638, 'gamma': 'auto'} | Fractals |
| | SVM | {'C': 37.55401188473625, 'gamma': 'scale'} | Binned |
| | SVM | {'C': 37.55401188473625, 'gamma': 'scale'} | Merged |
| **FOLD_4** | RandomForest | {'max_depth': 44, 'n_estimators': 120} | Fractals |
| | RandomForest | {'max_depth': 35, 'n_estimators': 314} | Binned |

| | | | |
|---|---|---|---|
| | RandomForest | {'max_depth': 44, 'n_estimators': 393} | Merged |
| | LightGBM | {'learning_rate': 0.18198808134726413, 'n_estimators': 234, 'num_leaves': 50} | Fractals |
| | LightGBM | {'learning_rate': 0.12973169683940733, 'n_estimators': 202, 'num_leaves': 48} | Binned |
| | LightGBM | {'learning_rate': 0.0849080237694725, 'n_estimators': 370, 'num_leaves': 37} | Merged |
| | SVM | {'C': 18.44347898661638, 'gamma': 'auto'} | Fractals |
| | SVM | {'C': 18.44347898661638, 'gamma': 'auto'} | Binned |
| | SVM | {'C': 37.55401188473625, 'gamma': 'scale'} | Merged |

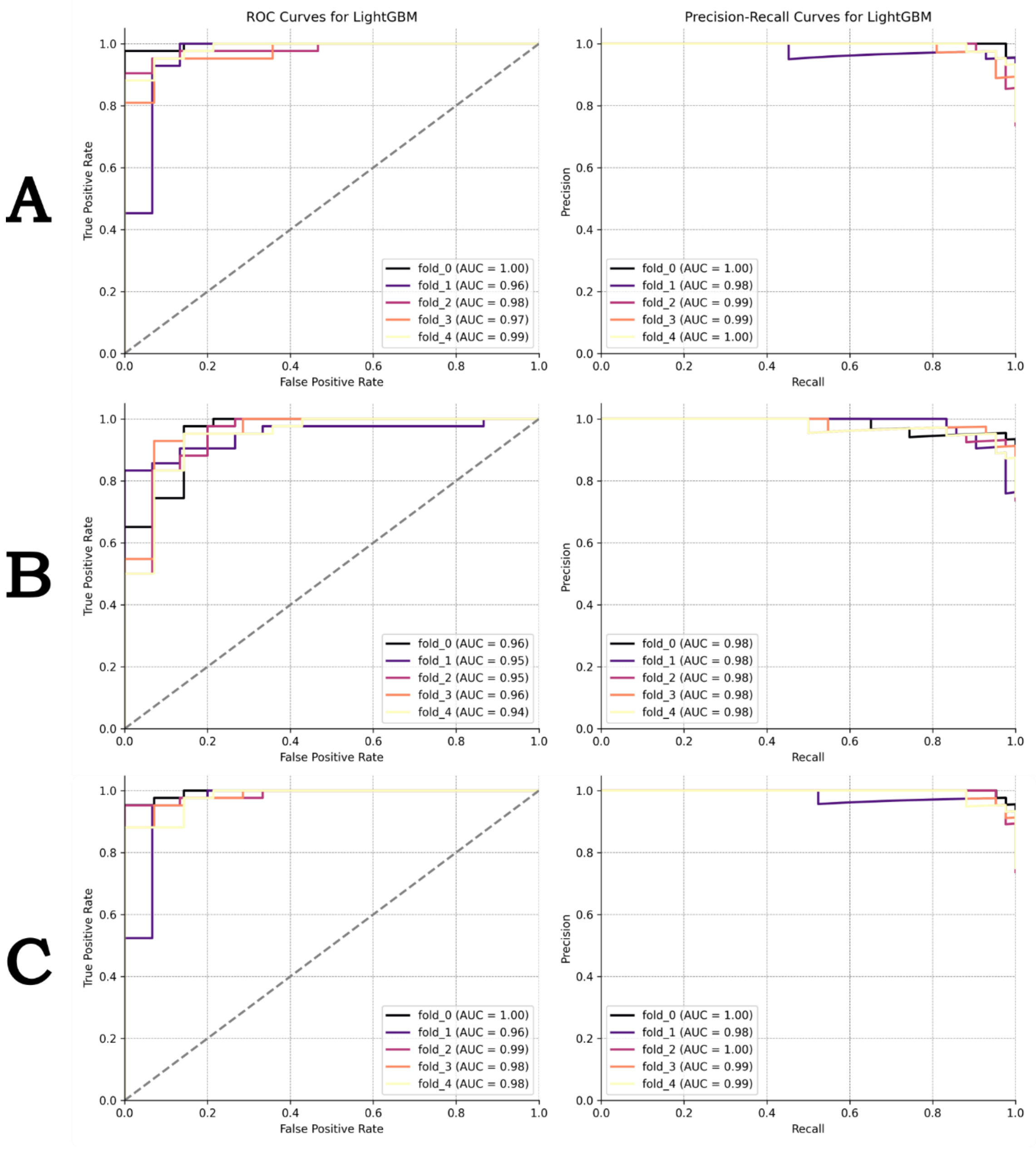


**Figure 1:** ROC and PR curves for each fold and each category tested for the LightGBM model. (A) Binned spectra. (B) Fractals features. (C) Merged features.

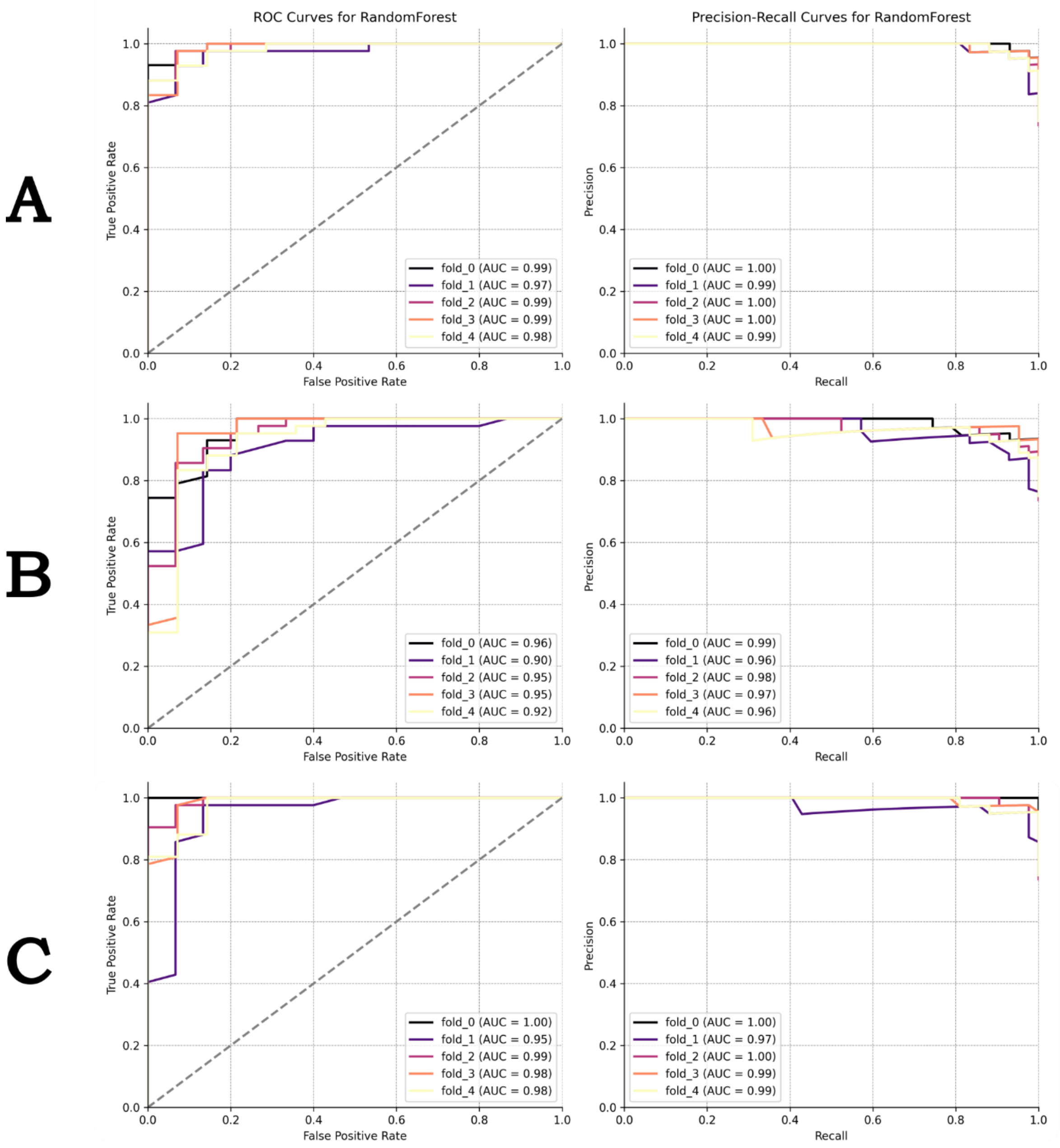


**Figure 2:** ROC and PR curves for each fold and each category tested for the Random Forest model. (A) Binned spectra. (B) Fractals features. (C) Merged features.

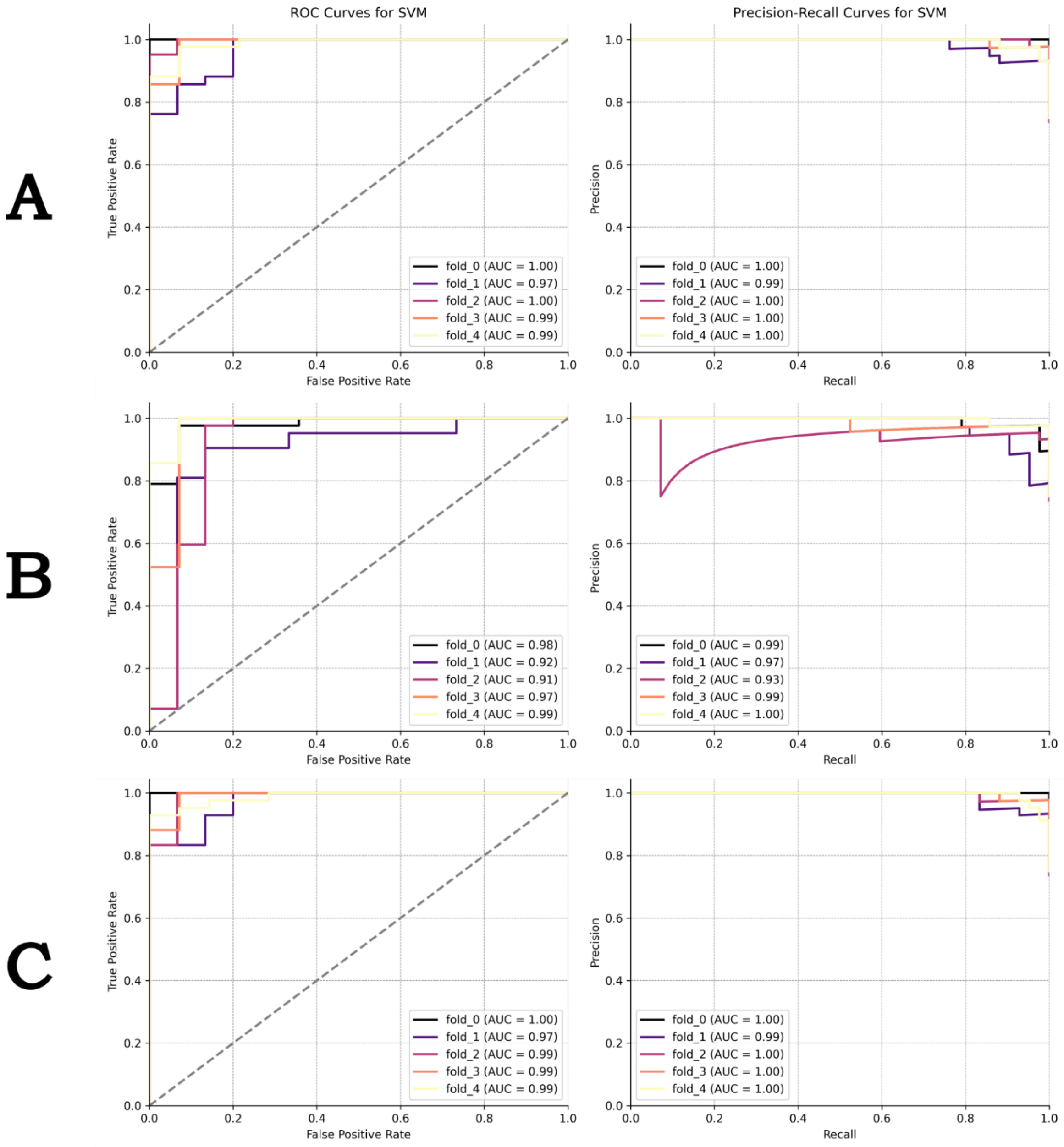


**Figure 3:** ROC and PR curves for each fold and each category tested for the SVM model. (A) Binned spectra. (B) Fractals features. (C) Merged features.

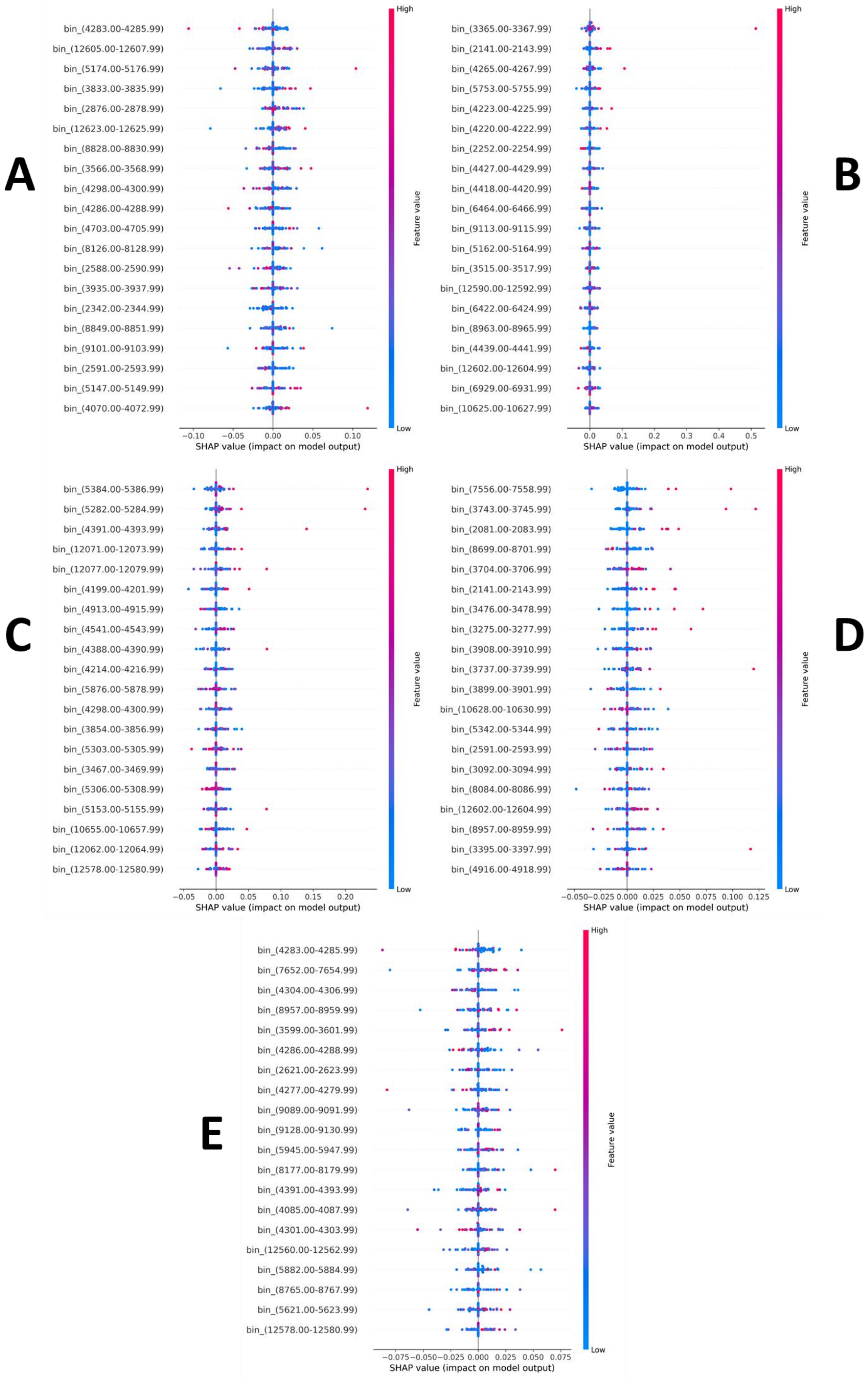


**Figure 4:** SHAP results for the binned dataset for each fold, starting at fold 1 (A) up to fold 5 (E).

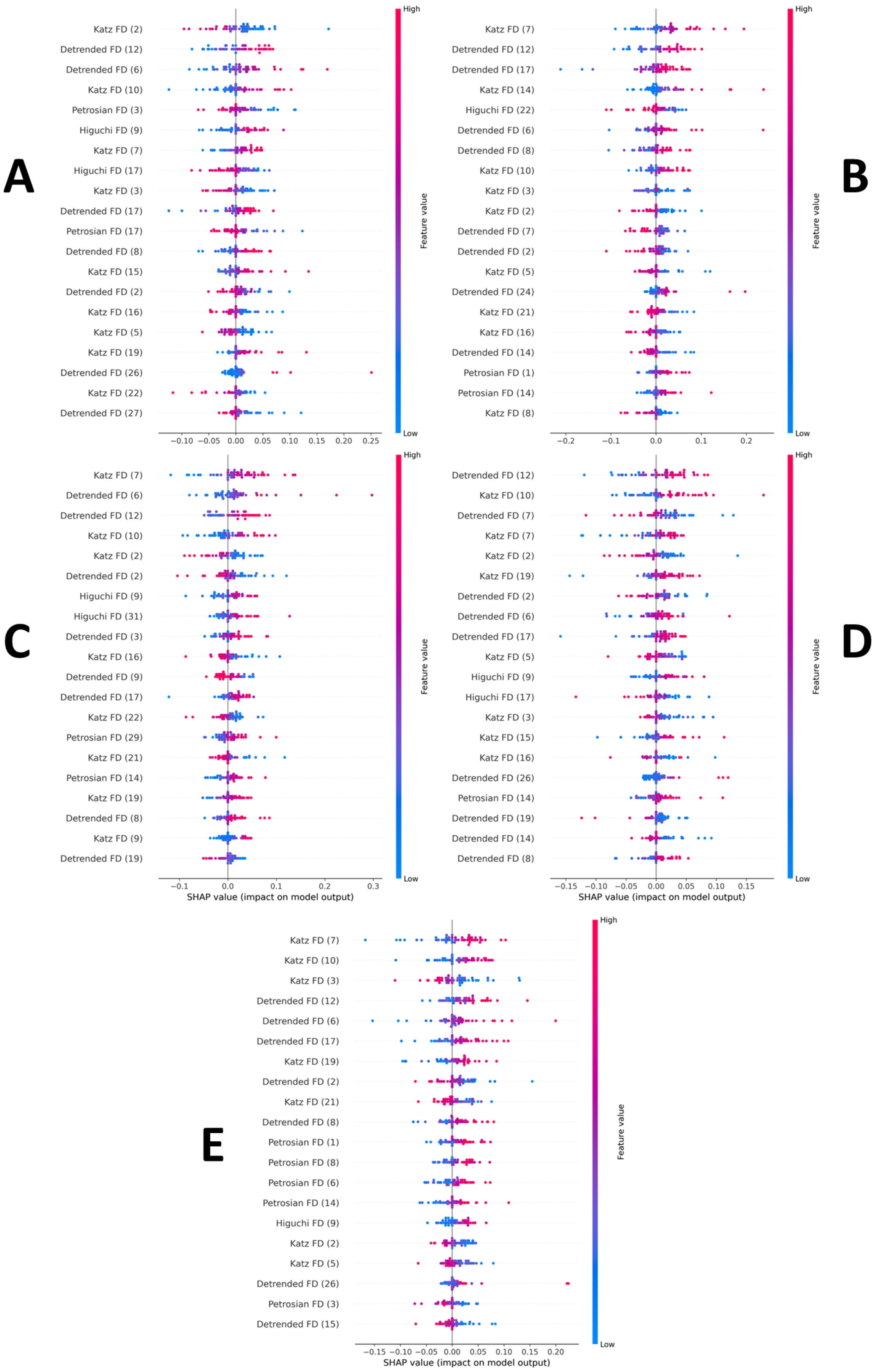


**Figure 4:** SHAP results for the fractals dataset for each fold, starting at fold 1 (A) up to fold 5 (E).
